# Density-dependent selection during range expansion affects expansion load in life-history traits

**DOI:** 10.1101/2022.11.08.515702

**Authors:** Mackenzie Urquhart-Cronish, Amy L. Angert, Sarah P. Otto, Ailene MacPherson

## Abstract

Models of range expansion have independently explored fitness consequences of life-history trait evolution and increased rates of genetic drift—or “allele surfing”—during spatial spread, but no previous model has examined the interactions between these two processes. Here, we explore an ecologically complex range expansion scenario that combines density-dependent selection with allele surfing, using spatially explicit simulations to asses the genetic and fitness consequences of density-dependent selection on the evolution of life-history traits. We demonstrate that density-dependent selection on the range edge acts differently depending on the life-history trait and can either diminish or enhance allele surfing. Specifically, we show that selection on the range edge is always weaker at sites affecting competitive ability (*K*-selected traits) than intrinsic growth rate ((*r*-selected traits). We then link differences in the frequency of deleterious mutations to differences in the efficacy of selection and rate of mutation accumulation across distinct life-history traits. Finally, we demonstrate the fitness consequences of accumulated deleterious mutations for different life-history traits are related to the population density in which they are expressed. Our work highlights the complex relationship between ecology and expressed genetic load, which will be important to consider when interpreting both experimental and field studies of range expansion.

## Introduction

Recent advances in evolutionary theory have highlighted the interaction between ecological and evolutionary processes during range expansion (Excoffier et al. 2009; Miller et al. 2020). These spatial eco-evolutionary dynamics are shaped by repeated population bottlenecks due to sequential founder events, which create a decreasing cline in population density from the range core towards the range edge. These spatial patterns in population density and effective population size impact the relative strength of selection on the range edge compared to the range core, even in the absence of underlying environmental gradients (Hallatschek et al. 2007). At low population densities, genetic drift can overwhelm selection and lead to the stochastic increase in frequency, and even fixation, of rare alleles (Edmonds et al. 2004). This process, referred to as ‘allele-surfing’ (first described by Eswaran 2002, and named by Edmonds et al. 2004; Klopfstein et al. 2006), can occur with beneficial, neutral, and deleterious mutations (Travis et al. 2007) and can have negative fitness consequences if deleterious alleles surf to high frequencies in range edge populations (i.e., expansion load) (Peischl et al. 2013).

Previous models of allele surfing have, however, ignored the ecological context of selection by assuming constant selection regardless of the current population size. This assumption is ecologically limited given that alleles affecting birth/death rates or competitive abilities are expected to experience different selection at low vs. high population densities. In particular, we expect selection during range expansion to reflect ecological conditions (e.g., population size differences) that are substantially different on the range edge than in the range core resulting in different strengths of intraspecific competition. The concept of r-versus K-selection (MacArthur and Wilson 1967) reflects this view that selection on genes will differ depending on their impacts on growth versus competition. Despite this, most previous work on allele surfing during range expansion has assumed constant selection, regardless of the competitive environment (Peischl and Excoffier 2015, Peischl et al. 2013 Peischl et al. 2015, Gilbert et al. 2017, Peischl and Gilbert 2020).

In empirical studies, expansion load is measured as a relative comparison between the fitnesses of core and edge individuals (Willi et al. 2018). Many field studies of expansion load have only assessed putative genetic signatures of load (e.g., *dN/ds* ratio; dePedro et al. 2021, Gonzalez-Martinez et al. 2017), and not the direct fitness consequences (e.g., heterosis). Drawing conclusions about expansion load from only genetic data is difficult due to the complex relationship between genotype, phenotype, and fitness (Svensson and Calsbeek 2012), the context-dependent nature of deleterious mutations (reviewed in Anderson et al. 2013), and the fact that putative genetic signatures of load can be confounded by demographic history and influenced by the presence of positive selection (Excoffier and Ray 2008, Brandvain and Wright 2016). Of the lab and field studies that investigated genetic signatures and fitness consequences of expansion load (Bosshard et al. 2017; Willi et al. 2018; Koski et al. 2019, Perrier et al. 2020), not all studies detected a direct relationship between genetic signatures and fitness consequences of expansion load (Takou et al. 2021).

Here, we examine the impact of density-dependent selection on life-history traits during range expansion. We focus on life-history evolution as these traits inherently differ in the presence of density dependence in classic demographic models (Roughgarden 1971). Although existing models of range expansion have investigated the effects of genetic drift (Polechová and Barton 2015; Polechova 2018) or density-dependent selection (Burton et al. 2010; Erm and Phillips 2020), to our knowledge, models of life-history evolution during range expansion have yet to include both density-dependent selection and allele surfing dynamics.

We evaluate the influence of density-dependent selection and allele surfing on both the genetic signatures and fitness consequences of life-history trait evolution during range expansion. As the eco-evolutionary dynamics leading to the accumulation of expansion load are potentially highly density-dependent, we explore dynamics of selection acting on three different life-history traits: birth-rate, competitive ability, and viability. We focus on cases where birth-rate and competitive-ability, but not viability, are density-dependent. We hypothesize that *1*) density-dependent selection on the range edge will act to diminish or enhance allele surfing relative to density-independent selection, and 2) fitness effects of genetic load will be related to the relative amount of allele surfing that occurs at the loci underlying each life-history trait. We first use a simplified single-population analytical model to derive expressions for selection coefficients as a function of population density. Then, we use simulation models to quantify genetic load following allele surfing, interpreting our results with the use of the selection coefficients. Our results have implications for how and when distinct life-history traits will accumulate expansion load and provide a more nuanced outlook for how the accumulation of expansion load may affect the fitness of edge populations in nature.

## Models & Results

### General description of the model

We developed a discrete-time individual-based model with overlapping generations in which individuals were tracked as they dispersed and reproduced across an environmentally uniform, one-dimensional landscape of demes. The life-cycle proceeded as shown in Fig. 1.

**Figure 1:**
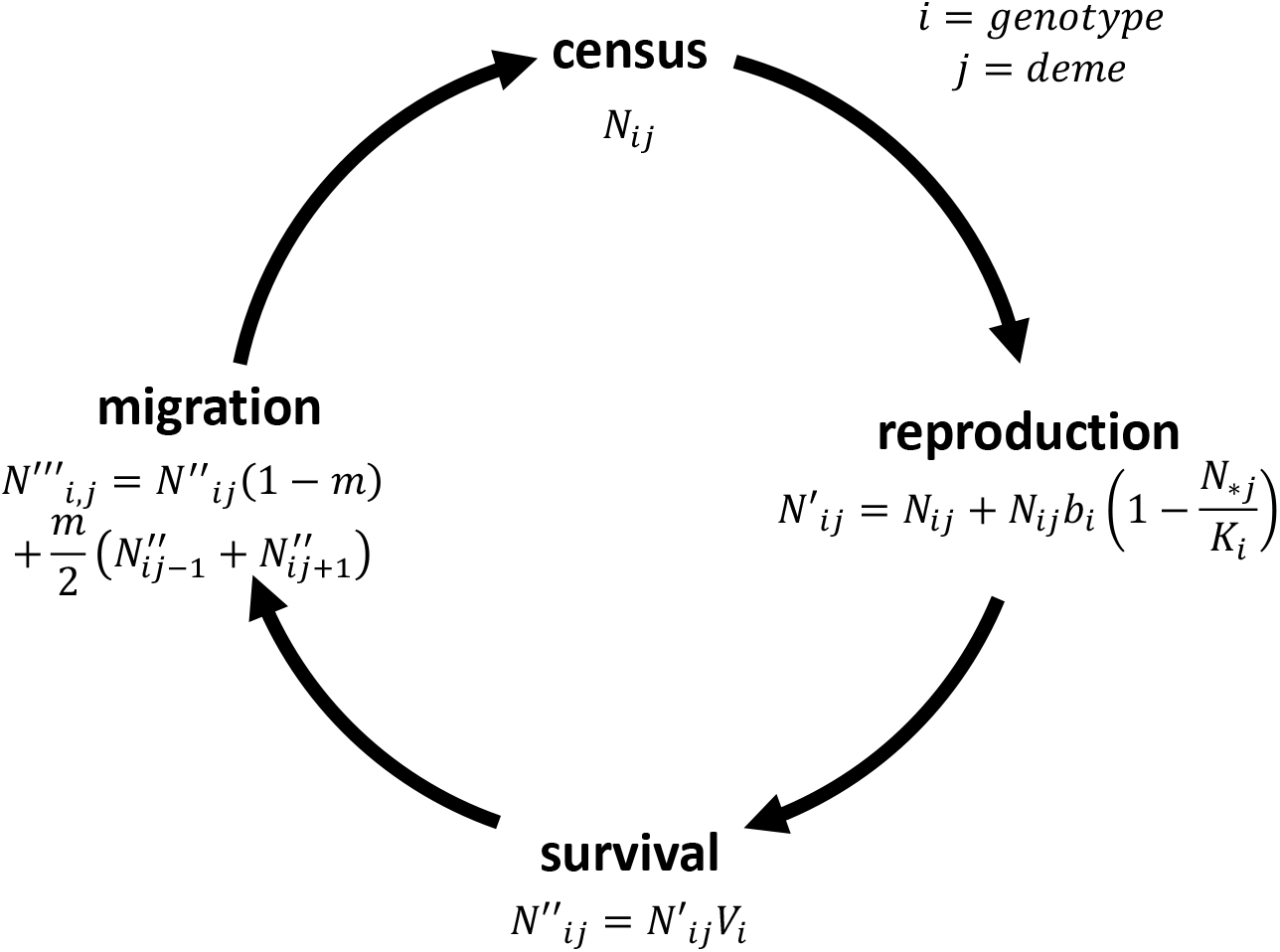
Life cycle of our model, with four distinct stages (census, reproduction, survival, and migration).

Selection occurs twice in the life-cycle, once during reproduction (fecundity selection) and again during survival (viability selection). When simulating surfing, individuals possess a genome of 300 biallelic codominant loci, with 100 loci each contributing to each of three life-history traits: birth-rate, viability, and competitive ability (Table 1). Loci underlying each trait are arranged randomly along the genome. Initially, all loci were homozygous wild-type with deleterious alleles arising by mutation, with a constant per-site mutation rate (*μ* = 0.0001) and free recombination (*r* = 0.5). We focus on the simplified case where there are no back mutations (mutant to wild-type mutation) to allow us to more easily detect deleterious mutation accumulation over time. Individuals mate randomly, including potential selfing, within their current deme. Individual fecundity is determined by an individual’s birth rate (*b_i_*) and competitive ability (*α_i_*), expressed via its inverse:

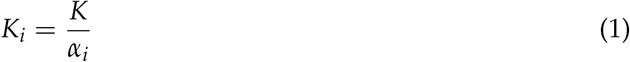

where *K* is proportional to the equilibrium deme size and sets the scale of the system. Following reproduction an individual with genotype *i* survives according to its viability, *V_i_*. The resulting expression for the absolute fitness of genotype *i* in a deme of total size *N*_*j_ is given by:

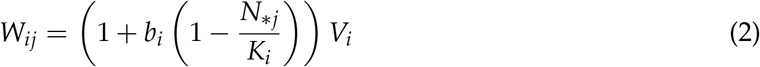

**Table 1:** Simulation model parameters.

| Parameters | Value |
| --- | --- |
| Number of generations burnin | 10 000 |
| Number of generations expansion | 5000 |
| Number of burnin demes | 5 |
| Number of expansion demes | 300 |
| Migration rate ($m$) | 0.005 |
| Number of loci (nLoci) | 300 |
| Mutation rate ($\mu$) | 0.0001 |
| Recombination rate ($r$) | 0.5 |
| Phenotypic effect ($x$) | 0.01 |
| Baseline birth-rate ($b_0$) | 1.1 |
| Baseline viability ($V_0$) | 0.7 |
| Baseline carrying capacity ($K_0$) | 125 |

The value of each life-history trait of an individual with genotype *i, z_i_* = {*V_i_, b_i_, K_i_*} is determined by the multiplicative effects of the underlying codominant loci:

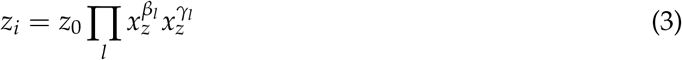

The genotype, *i*, in the above equation is specified by the indicator variables *β_l_, γ_l_*, which take on values of 0 or 1 if the genotype carries the wild-type or mutant allele at locus *l* on the maternal (parental) chromosome respectively. The constant *z*_0_ determines the baseline phenotype of the initial wild-type genotype (*β_l_ = γ_l_ =* 0 for all *l*). For the three focal life-history traits these baseline parameter values are given by *b*_0_, *V*_0_, and *K*_0_ (Table 1) and *x_z_* is a constant with a value less than or equal to one that determines the fitness component value in the presence of a deleterious mutant allele.

From (Equation 2) describing the fitness of each genotype given the current population size, we can obtain an expression for the selection coefficient (*s*_z_ (*N*_*j_)) for each trait (Table 2) for a given phenotypic effect of mutation (*x*_z_) for these three traits. Importantly here, the selection coefficients (*s*_b_, *s*_K_, but not *s*_V_) are functions of the total population size of the current deme *N*_*j_, indicating that selection is density-dependent. While the realized strength of selection varies with population density, we can equalize selection on each trait by defining the mutational effects of each trait *x*_z_ such that the selection coefficients are equal (*s*_b_ = *s*_V_ = *s*_K_) when the population size is at the ecological equilibrium:

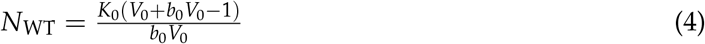

**Table 2:** Selection coefficients and phenotypic effects of mutations on each fitness component in our analytical model. Equations are for a wild-type population at its ecological equilibrium, where the efficacy of selection is equivalent across the three distinct life-history traits.

| Life-history trait | Selection coefficient | Phenotypic effect of mutation |
| --- | --- | --- |
| Birth-rate | $s_b = \frac{b_0(K_0 - N_{WT})(1 - x_b)}{K_0 + b_0 K_0 - b_0 N_{WT}} = x - 1$ | $x_b = \frac{V_0 - x}{V_0 - 1}$ |
| Viability | $s_V = x - 1$ | $x_V = x$ |
| Competitive ability | $s_K = \frac{b_0 N_{WT}(1 - x_K)}{K_0 + b_0 K_0 - b_0 N_{WT}} = x - 1$ | $x_K = \frac{-1 + V_0 + b_0 V_0}{V_0 + b_0 V_0 - x}$ |

(Table 2, see *Mathematica* notebook for derivation).

After selection (reproduction and survival), individuals migrate with a probability *m* to one of the two closest demes (Table 1). We initially investigated three different dispersal modes (stepping-stone, geometric dispersal, rare long distance dispersal) and did not detect any substantial differences in simulated range expansion dynamics (Fig. S1), so we only present results with a stepping-stone model. To analyze the dynamics of genetic drift during range expansion we calculate two quantities: the allele frequency and expansion load near the range edge. Because population size on the range edge (the most recently colonized deme) is often very small and highly stochastic, we measured the dynamics of allele frequencies (*p_edge_*), phenotypic expression of life-history traits (*z_edge_*) (Equation 3), and fitness (*W_edge_*) one deme behind the edge.

An important consequence of density-dependent fitness is that expansion load no longer has a simple definition but is instead dependent on the ecological context:

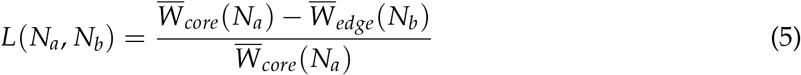

Expansion load is defined as the change in fitness due to genetic differences in the populations caused by range expansion. If fitness differences are measured at different population densities (i.e., (*N_a_* ≠ *N_b_*), then the resulting measure is a biased estimate of load.

In a density-dependent context, fitness depends on population size (Equation 5) where the over-bars indicate the average fitness of individuals within the deme indicated in the subscript. As a result, load is a function of the two population sizes *N_a_* and *N_b_* in which the core and edge fitnesses are measured. To illustrate how load depends both qualitatively and quantitatively on population size we contrast two scenarios: where both fitnesses are measured experimentally using the edge population density (*N_a_* = *N_b_* = *N_edge_*) and both core densities (*N_a_* = *N_b_* = *N_core_*). We also examine a scenario where fitness is calculated using local densities (*N_a_* = *N_core_* and *N_b_* = *N_edge_*), and this scenario leads to a biased estimate of load since fitness is not measured across the same environment. The experimental design used to detect load must match the first two scenarios, as the final scenario with local densities does not meet the definition of expansion load. Finally, to partition load into the contributions from each life-history trait, we find the mean fitness by averaging over the variation in a focal trait within the deme of interest while replacing the remaining two fitness components with their respective baseline values (*b*_0_, *V*_0_, or *K*_0_) in Equation 2.

The key point is that deleterious alleles primarily affecting competition (teal Fig. 2A) experience weaker selection in smaller populations and should show elevated surfing, while those affecting birth-rate are more strongly disfavoured in smaller populations (purple Fig. 2A) and so should accumulate less often at the range edge. By contrast, alleles affecting viability in a density-independent manner experience the same strength of selection regardless of population size (yellow Fig. 2A), as previously assumed in models of genetic surfing (Peischl et al. 2013, Peischl et al. 2015)

**Figure 2:**
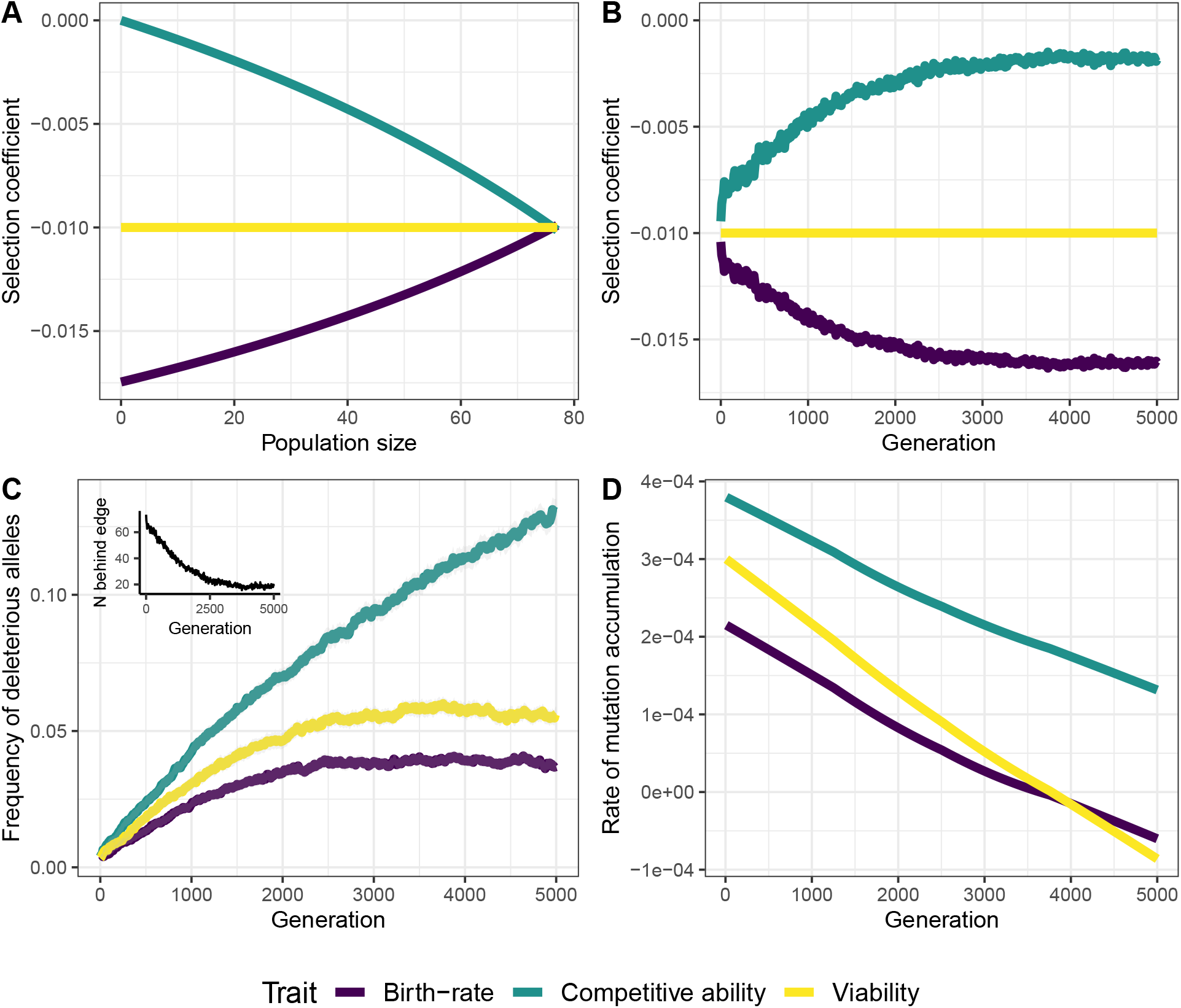
Population size during range expansion affects selection. (A) Fitness model for how selection coefficient (Table 2) changes with population size (x-axis: small to large, left to right). (B) Selection coefficients observed in simulations on the range edge (x-axis is opposite direction of panel A: population size decreases left to right over time). Inset shows changes in population size on the range edge during range expansion. (C) Frequency of deleterious mutations that accumulated at loci underlying distinct life-history traits during range expansion. (D) The rate of mutation accumulation over time, calculated from the slope of a loess model fit to the deleterious allele frequency curves in panel B.

Individual-based simulations were coded in C++ and run on the Compute Canada cluster. Analyses of simulation results were conducted in R (Version 4.0.3, R Core Team 2020) using R Studio software (version 2021.09.2). The model code and simulation outputs can be found here (https://www.dropbox.com/sh/cjklynqtutzgiqv/AAD3YaiGDddN1ePqvh-RzC3Qa?dl=0, will be uploaded to Dryad at time of publication).

### Individual-based simulations

Populations were initiated at a carrying capacity of 76 individuals per deme in the leftmost five demes (i.e., range core) of our one-dimensional 300 deme landscape. After a burnin period of 10000 generations where populations reached eco-evolutionary equilibrium, the remaining demes in the landscape became available, allowing for range expansion to occur. Expansion occurred over 5000 generations. Simulations were run in parallel, and 50 replicates were conducted per burnin and expansion simulation run. We also investigated a scenario where the phenotypic effects of mutations were identical (Fig. S2, i.e., *x_b_* = *x_K_* = *x_V_* = *x* relative to values in the main text, see column 3 of Table 2) and found similar results.

### Density-dependent selection during range expansion depends on life-history trait

Population density on the range edge decreases over time (inset Fig. 2C) facilitating the accumulation of deleterious mutations on the range edge at loci underlying all life-history traits, as captured by the genome-wide average allele frequencies (Fig. 2C). The extent of allele surfing and the number of deleterious alleles that accumulated during range expansion differed among our three life-history traits. As expected, loci underlying birth-rate accumulated the fewest deleterious alleles over time, while loci underlying competitive ability accumulated the greatest number.

Because population size declines with range expansion (inset Fig. 2C) selection on loci underlying birth-rate becomes stronger over time, whereas selection on competitive ability relaxes over time (Fig. 2B). By contrast, the strength of selection on viability remains the same, as it is density independent (Fig. 2A & B). Figure 2D shows the rate of mutation accumulation at loci underlying competitive ability remains consistently higher than the rate of mutation accumulation in birth-rate or viability, as expected from differences in the strength of selection (Fig. 2C). Note that the rate of mutation accumulation slows (Fig. 2D), eventually reversing as migration of beneficial alleles from the core displace deleterious alleles that have surfed at the range edge (observed around generation 4000 for birth-rate and viability).

### Expansion load varies among life-history traits

The observed expansion load depended on the population density at which fitness was measured, demonstrating that expansion load depends on the experimental design. When a single population density (either core or edge) is used to calculate fitness, the accumulation of deleterious mutations during allele surfing results in expansion load, as expected (Fig. 3). If local densities are used, however, this results in a biased estimate of load where values of genetic load are negative (Fig. 3). This is because expansion load is defined by genetic change in a common environment, and using local population densities introduces the possibility of bias (see second term in Equation 6).

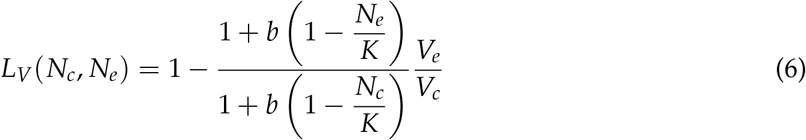

**Figure 3:**
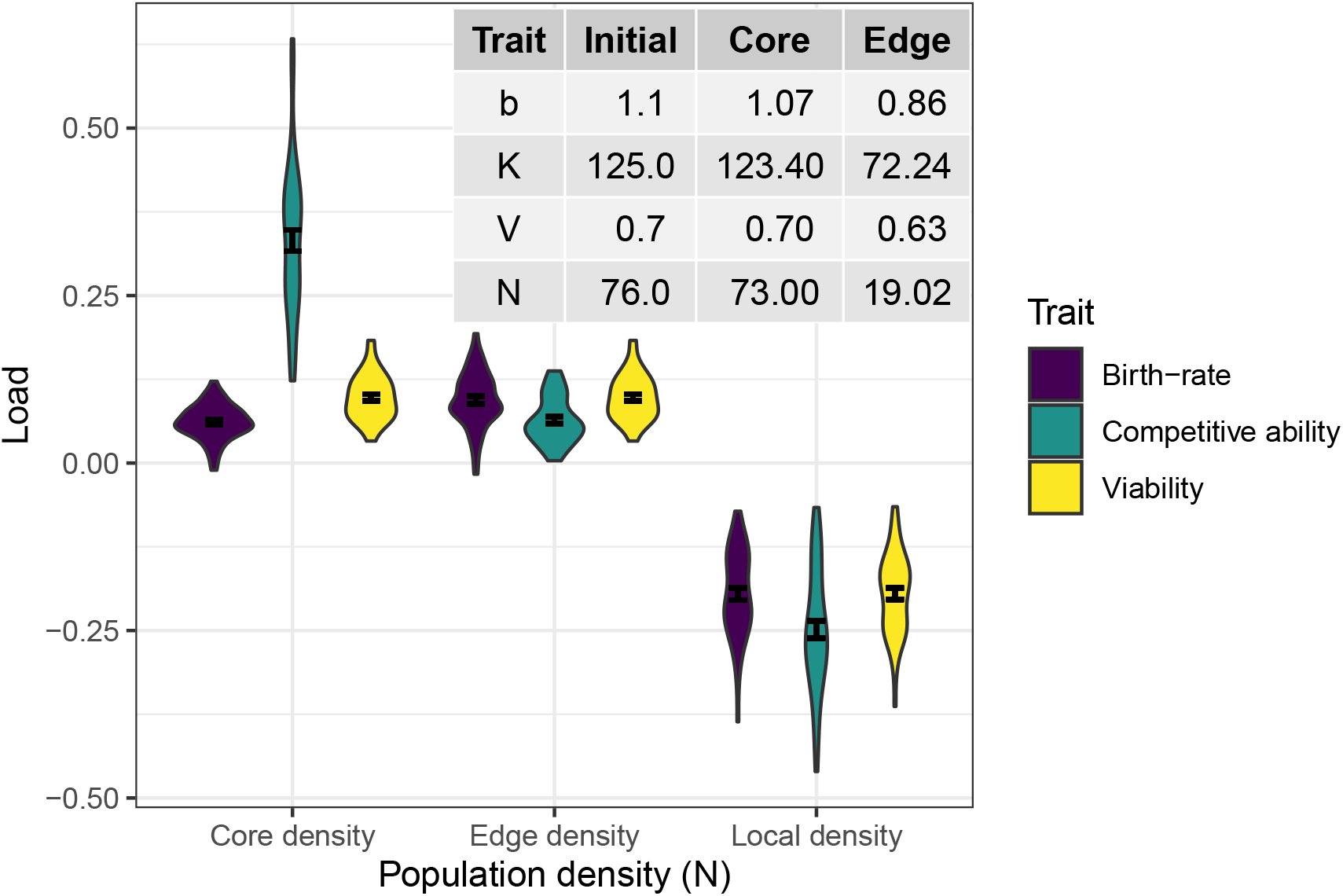
Expressed expansion load depends on experimental population densities where fitness is measured. Violin plots, along with the average expansion load (± standard error) for different life-history traits calculated using average population density at generation 5000 after expansion began in the range core (Core density) and at the range edge (Edge density). At local population densities (Local density) a biased estimate of load is measured since the environment where fitness is measured is not uniform. To calculate fitness, each focal life-history trait was calculated using the average fitness component value at the end of range expansion with the other two traits held at their initial values (inset Table; b = birth-rate, K = competitive ability, V = viability, N = population density). The inset table also shows the average value of life-history traits and population sizes used to calculate fitness in the core and at the edge (Equation 2).

The relative contribution of each life-history trait also changes with the population density used in an experiment. Namely, mutations in loci underlying competitive ability contributed most to the expansion load when measured at population densities typical of the core and contributed less to expansion load when measured at densities typical of the range edge (Fig. 3). Although fewer mutations accumulated at loci underlying birth rate during range expansion (Fig. 2C), those mutations made a relatively large contribution to expansion load in the edge scenario. As expected the expressed load in density-independent viability remained constant when measured either at the core or edge density.

## Discussion

We demonstrated that density-dependent selection during range expansion interacts with allele surfing on the range edge. While stronger genetic drift is always expected on the range edge, strong selection on loci underlying traits that improve population performance at low population densities can attenuate allele surfing in that portion of the genome. According to our results, surfing should disproportionately affect loci underlying competitive ability (*K*-selected traits), which experience relaxed selection on the range edge, but should have more limited impacts on *r*-selected traits such as birth-rate, with intermediate impacts for density-independent traits (in our study, survival). It is important to note that these fitness measures were taken at generation 5000 after the opening up of the range, when populations at the range edge were still small (inset, Figure 2C). Once range expansion is over, if population sizes equilibrate at similar levels across the range, we would expect the expansion load (while it remains) to mimic the patterns observed at the core population density (Figure 3 left). Importantly, these results demonstrate the experimental population densities used and the life-history traits assessed influence the value of the expansion load that is measured.

Given the importance of environmental context (population density) for calculating fitness and expansion load, we reflect on previous empirical investigations of expansion load to interpret their results through the lens of density-dependent selection. Empirical studies of expansion load have mainly focused only on detecting the putative genetic signatures of load (Henn et al. 2016, Zhang et al. 2016; González-Martínez et al. 2017; dePedro et al. 2021). These studies are indirect and do not reveal the relationship between the extent of deleterious mutation accumulation and the expansion load (more mutations does not always equal more load, as we find with competitive ability observed at population densities at the range edge). In order to make informative inferences about expansion load, it is important to directly link genetic signatures of deleterious mutations to measures of fitness (Bosshard et al. 2017; Willi et al. 2018; Koski et al. 2019; Takou et al. 2021), and our results have demonstrated that measuring fitness only using a single population density may not be sufficient since the population density where fitness is measured has a large influence on the observed expansion load. Some studies measure fitness in a common garden (Orsucci et al. 2020, Willi et al. 2018, Perrier et al. 2020), but none have measured fitness under varied intraspecific population densities. As a practical extension of our simulation study, we suggest empirical studies of expansion load measure fitness at distinct population densities to capture this dependence.

### Future directions to model density-dependent selection on life-history traits during range expansion

We designed our eco-evolutionary model to detect expansion load which means we specifically set parameter values with an intermediate migration rate (so that colonization of adjacent demes would occur before equilibrium was reached but not so fast that there was panmixia) and no underlying environmental gradient (Peischl et al. 2013). Among our many simplified assumptions, we did not allow for the evolution of dispersal ability (Peischl and Gilbert 2020), life-history trade-offs (Burton et al. 2010), or variation in environmental conditions across the range (Gilbert et al. 2017). Below, we discuss the implications of our assumptions and what more complex models that incorporate these caveats may or may not show us about how density-dependent selection on the range edge affects the accumulation of expansion load.

While we investigated and found no substantive effect of different dispersal modes on range expansion in our simulations (Fig. S1), we did not explicitly allow for the evolution of dispersal ability or density-dependent dispersal in our model. Both evolution of greater dispersal ability (via spatial sorting, Shine et al. 2011) and density-dependent dispersal are expected to reduce the overall expansion load (Peischl and Gilbert 2020). Greater dispersal ability will result in faster range expansion with fewer generations for allele surfing to occur and mutations to accumulate. Similarly, if dispersal rates increase with population density this would result in higher populations densities on the range edge and reduced effects of genetic drift and therefore allele surfing. However, it should be noted that populations can potentially evolve to overcome density-dependent dispersal that acts to slow down range expansion (Erm and Phillips 2020), approaching the case considered here where population density on the range edge is lower than the core.

In our model, range expansion occurred along a one-dimensional landscape with no underlying environmental gradient. Gilbert et al. (2017) investigated the effect of underlying environmental gradients during range expansion on the accumulation and fitness consequences of expansion load, finding that environmental gradients slow down range expansion by reducing colonization success of dispersing individuals due to a mismatch between colonizing individuals and edge habitat conditions. Depending on the steepness of the environmental gradient, this can create conditions where range expansion is stalled. This is a similar outcome to the density-dependent dispersal scenario where establishment is more difficult resulting in higher density on the range edge, which relieves expansion load. We would thus expect such gradients to reduce the difference in selection among life-history traits observed here.

Incorporating a two-dimensional landscape, we would expect deleterious mutations to accumulate at the same average rate as one-dimensional expansions (Peischl et al. 2013). However, we would also expect populations to recover more quickly from expansion load in two-dimensional expansions due to larger effective population sizes on the range edge making selection more efficient after expansion. Nevertheless, we would expect a decreasing population density from core to edge, so we would expect that competitive traits would still be more likely to surf and traits contributing to growth would be less likely to surf.

Investment in growth and reproduction often trade off, given that only a finite amount of resources are available to an individual (Stearns 1989). Previous models of range expansion have investigated co-evolution of density-independent fitness and dispersal in the context of expansion load (Peischl and Gilbert 2020) or used phenotypic models to investigate trade-offs among distinct life-history traits (Burton et al. 2010), but no current model incorporates allele surfing dynamics and density-dependent life-history trade-offs with an explicit genetic basis. Recent results demonstrate the evolution of enhanced reproductive ability of beetles at the range-edge, suggesting strong density-dependent selection during natural range expansion (Clark et al. 2022). Incorporating a life-history trade-off between birth-rate and competitive ability under density-dependent selection in a model of expansion load could reveal insights into the long-term effects of expansion load on these trade-offs.

### Conclusions

By incorporating density-dependent selection of life-history traits, our range expansion model provides a more nuanced understanding of existing expansion load (Peischl et al. 2013; Peischl and Excoffier 2015; Peischl et al. 2015; Gilbert et al. 2017; Peischl and Gilbert 2020). We demonstrated that the relationship between deleterious mutation accumulation underlying distinct life-history traits and the associated expressed expansion load is context-dependent, changing with the population density, both historically and at which fitness is measured. We further suggest suitable empirical systems (Willi et al. 2018; Orsucci et al. 2020) primed to test the implications of our model and results. By measuring expansion load at core population densities, such empirical work could examine if as predicted expansion load is exaggerated in traits under positive density-dependent selection (e.g., competitive ability) and suppressed in traits under negative density-dependent selection (e.g., birth rate).

## Acknowledgements

Feedback from M.C. Whitlock on an earlier version of the manuscript improved the final manuscript. M. Boehm and T. Usui provided helpful guidance writing R code. The authors were supported by NSERC (Discovery Grants to AA [RGPIN-2019-XXXX], AM [RGPIN-2022-XXXX], & SPO [RGPIN-2022-03726], CGS-D graduate scholarship to MUC), the University of British Columbia (Four-year Fellowship and Tuition funds to MUC), and the L’Oreal FCRF (MUC).

## Author contributions

MUC, AM, and SPO conceived of the study, with input from AA. MUC and AM designed the simulation model with input from SPO and AA. AM wrote the simulation code in C++ and the analytical model in Mathematica, MUC ran the simulations and wrote the R code for data processing, analysis, and figure generation. MUC wrote the first draft of the manuscript, with input and contributions from all authors.

## Data accessibility

dataacc All data and analysis code used in this article will be deposited in a repository (e.g., Dryad) following publication. Data during the review process can be accessed on Dropbox: https://www.dropbox.com/sh/ciklynqtutzgiqv/AAD3YaiGDddN1ePqvh-RzC3Qa?dl=0

## Supplementary Figures

**Figure S1:**
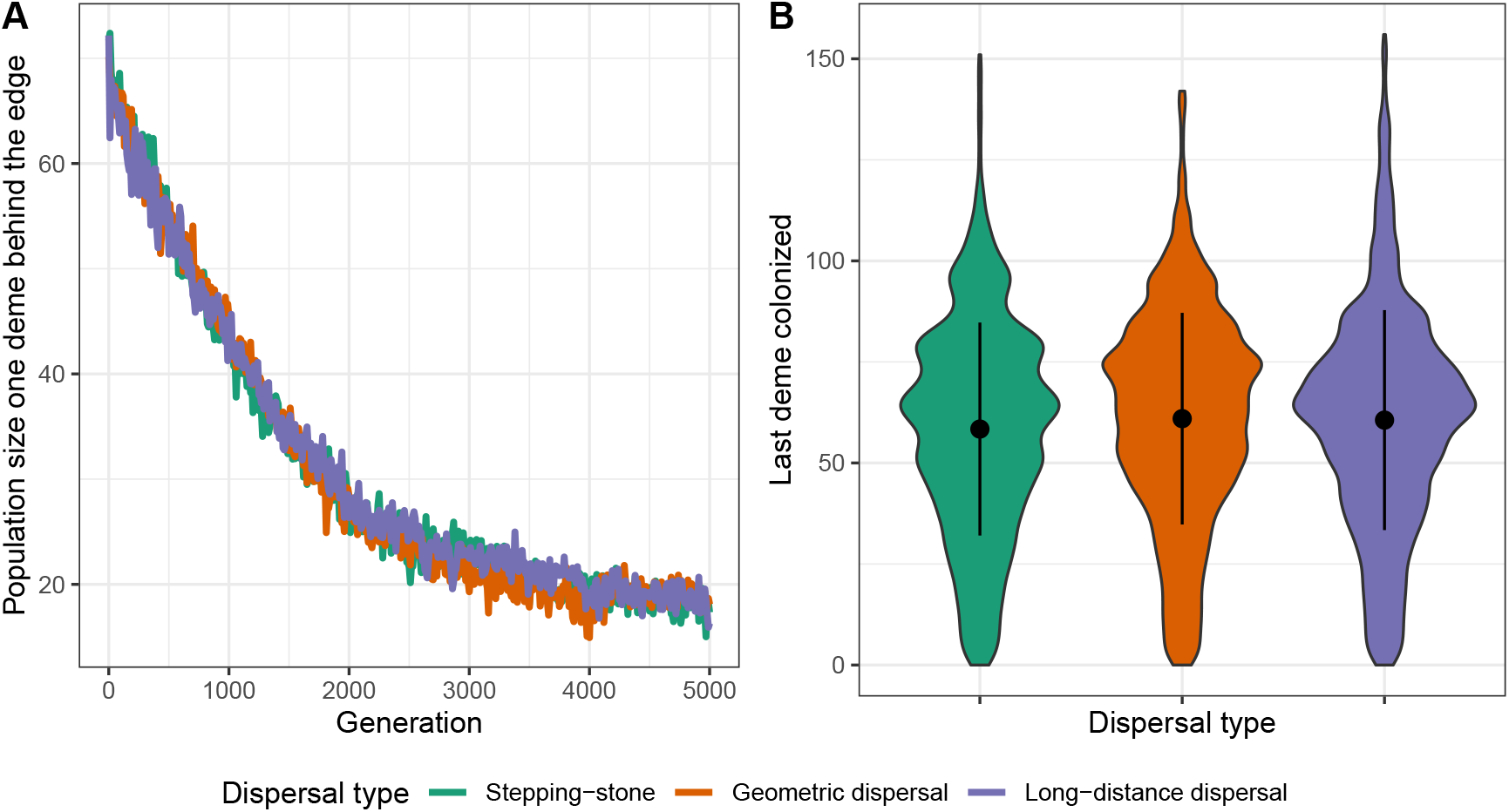
Ecological dynamics under three different dispersal scenarios. No differences in A) population size behind the range edge or B) final deme colonized across a comparison of simulated range expansions with different dispersal types (geometric dispersal, long-distance dispersal, stepping-stone). Stepping-stone dispersal captures migration to neighbouring demes only with a probability of moving either left or right of (*m*/2). Geometric dispersal refers to the case where the number of demes an individual moves either left or right (with probability of 0.5) is geometrically distributed with parameter (*p* = 1 – *m*). In Long-distance dispersal individuals move either left or right 1 deme with probability (*m* – *m*^2^/2) and a random number of demes between 1 and 5 with probability (*m*^2^/2). Note that the mean number of demes moved in each case is approximately equal.

**Figure S2:**
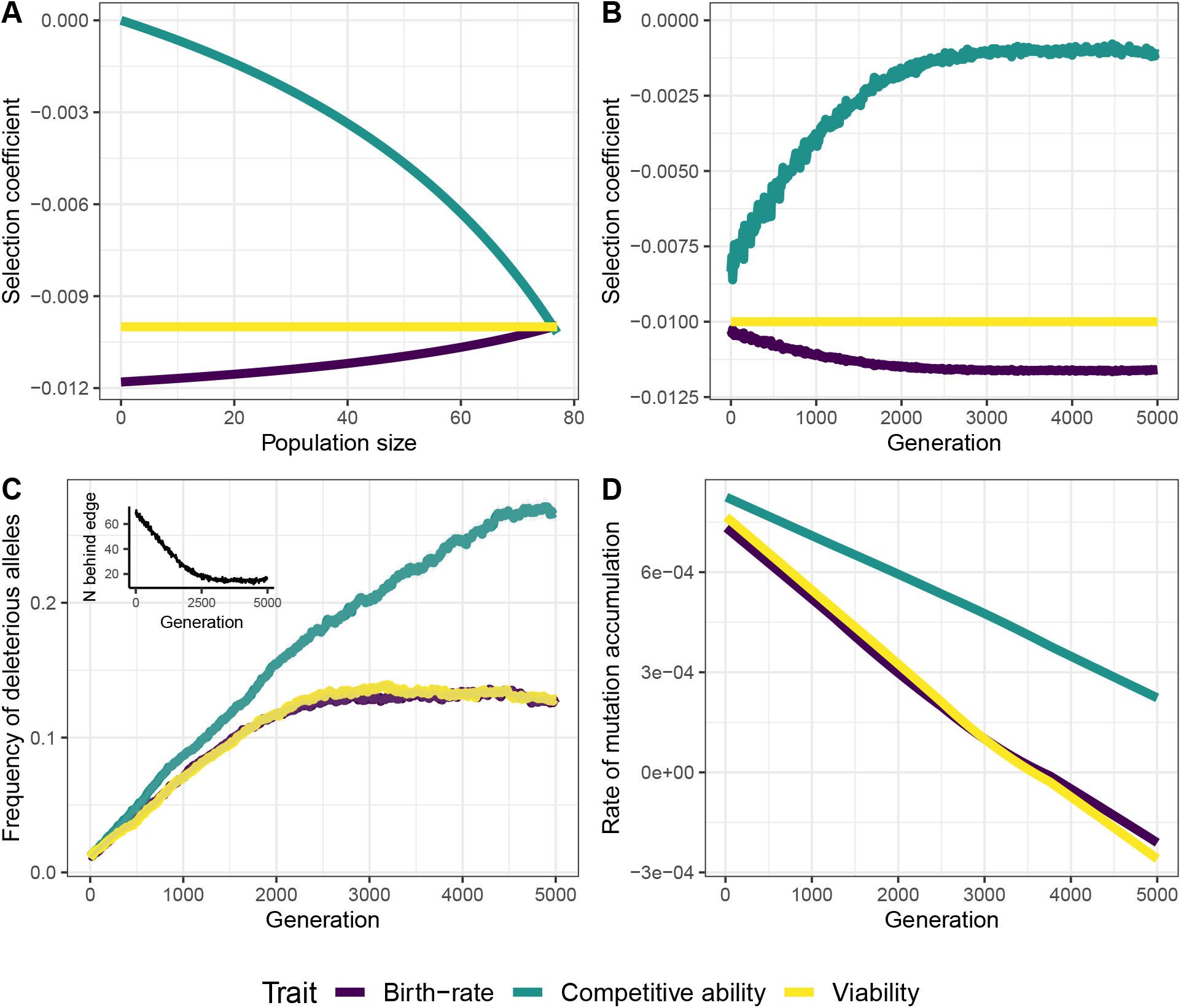
Figure is identical to Figure 2, but with different values of *b*_0_ = 7.7, *K*_0_ = 125, and *V*_0_ = 0.25 which is a scenario where the phenotypic effects of mutations were more similar (Equation 3) (A) Analytical model predictions for how the selection coefficient (see Table 2 for equations) changes with population size. (B) Frequency of deleterious mutations that accumulated at loci underlying distinct life-history traits during range expansion. Inset shows changes in population size on the range edge during range expansion. (C) Selection coefficients observed in simulations on the range edge. (D) The rate of mutation accumulation over time, calculated from the slope of a loess model fit to the deleterious allele frequency curves in panel B.

**Figure S3:**
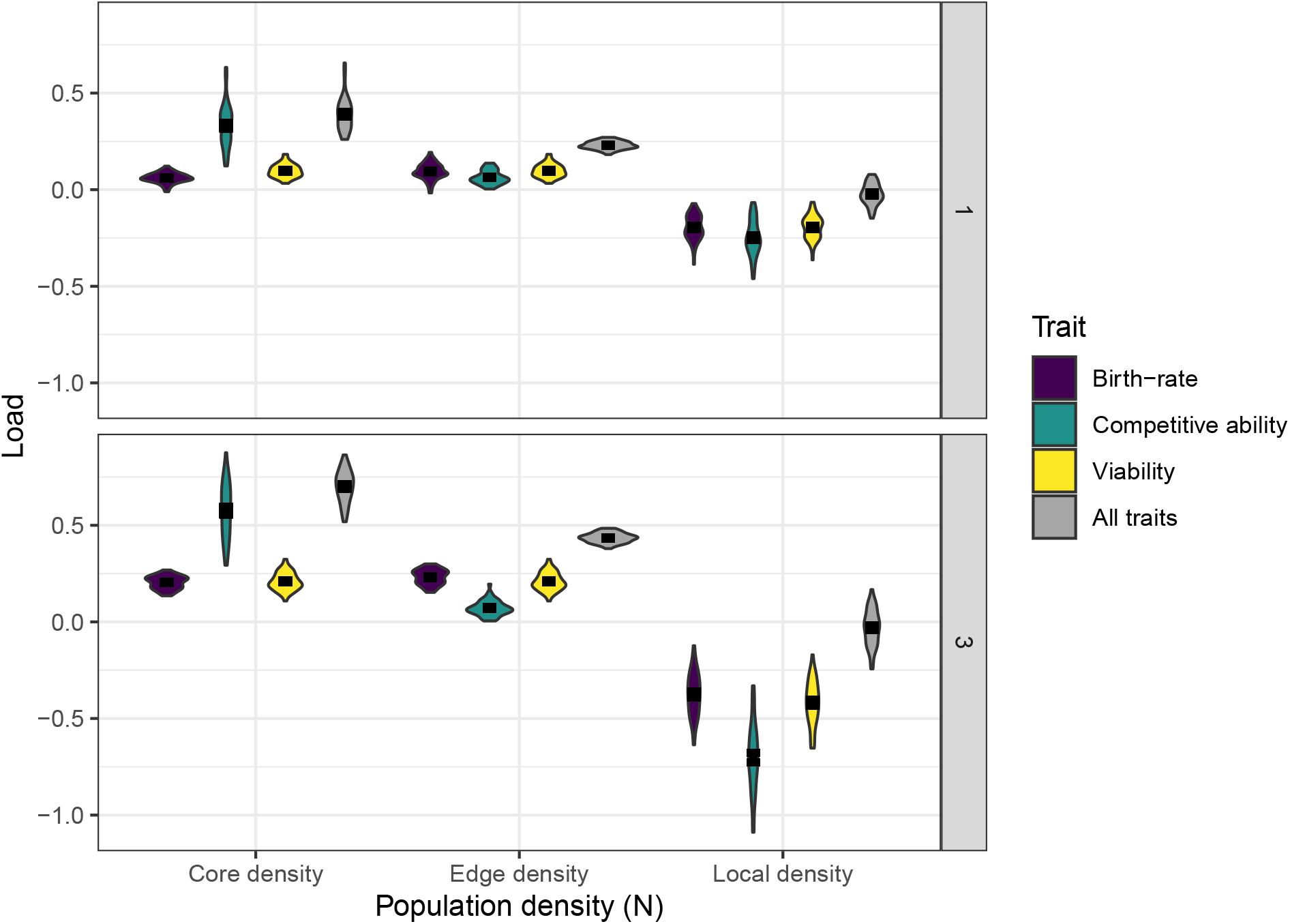
Violin plots of average replicate (n = 50) load plotted with total average expansion load (error bars ± standard error) for different life-history traits calculated using average population density in the range core (Core density)and at the range edge (Edge density). At local population densities (Local density) a biased estimate of load is measured since the environment where fitness is measured is not uniform. Panel 1 refers to parameter values in Table 1 and Panel 3 refers to parameter values used in Fig. S2. To calculate fitness, each focal life-history trait was calculated using the average fitness component value at the end of range expansion, while the other two traits were held at their initial values (inset Table; b = birth-rate, K = competitive ability, V = viability, N = population density). In the “All traits” scenario, all life-history traits used to calculate fitness based on their values at the end of range expansion.

